# Optimizing *E. coli* as a formatotrophic platform for bioproduction via the reductive glycine pathway

**DOI:** 10.1101/2022.08.23.504942

**Authors:** Seohyoung Kim, Néstor David Giraldo, Vittorio Rainaldi, Fabian Machens, Florent Collas, Armin Kubis, Frank Kensy, Arren Bar-Even, Steffen N. Lindner

## Abstract

Microbial C1 fixation has a vast potential to support a sustainable circular economy. Hence, several biotechnologically important microorganisms have been recently engineered for fixing C1 substrates. However, reports about C1-based bioproduction with these organisms are scarce. Here, we describe the optimization of a previously engineered formatotrophic *Escherichia coli* strain. Short-term adaptive laboratory evolution enhanced biomass yield and accelerated growth of formatotrophic *E. coli* to 3.3 g-CDW/mol-formate and 6 hours doubling time, respectively. Genome sequence analysis revealed that manipulation of acetate metabolism is the reason for better growth performance, verified by subsequent reverse engineering of the parental *E. coli* strain. Moreover, the improved strain is capable of growing to an OD_600_ of 22 in bioreactor fed-batch experiments, highlighting its potential use for industrial bioprocesses. Finally, demonstrating the strain’s potential to support a sustainable, formate-based bioeconomy, lactate production from formate and CO_2_ was engineered. The optimized strain generated 1.2 mM lactate—10 % of the theoretical maximum—providing the first proof-of-concept application of the reductive glycine pathway for bioproduction.

## Introduction

The valorization of carbon dioxide is a major challenge in our society and is subject to intense research and investment. Naturally, the biological transformation of carbon dioxide takes place in plants and algae on a massive scale. However, photosynthetic carbon fixation greatly suffers from low-energy conversion efficiencies in the range of 3–5% (Janssen et al., 2003; Zhu et al., 2010). Pure chemical transformation, where various chemicals such as urea, methanol, and salicylic acid can be derived directly from carbon dioxide, might be another option (He et al., 2013; Wong, 2014). However, such processes rely on extreme conditions and suffer from a limited product spectrum and low product selectivity. An emerging solution is to integrate biological and chemical processes to combine their individual strengths and circumvent their weaknesses. In such an approach, carbon dioxide can be reduced electrochemically, using a renewable energy source such as solar or wind, to various C1 compounds. Among them is formic acid, which can be produced with a very high faradaic efficiency and can be utilized as a feedstock for microorganisms (Chong et al., 2016; Han et al., 2012; Yishai et al., 2016). Formic acid is one of the most suitable C1 compounds for the bioindustry, especially because of its solubility and low toxicity (Claassens et al., 2019).

However, engineering natural C1-assimilating microorganisms to produce value-added biochemicals from single-carbon compounds is often limited by their poor growth characteristics and recalcitrance to genetic modification. Various natural C1 assimilation routes have been identified, including the reductive pentose phosphate cycle, the reductive acetyl-CoA pathway, and the reductive citric acid cycle from various domains of life (Bar-Even et al., 2012; Bar-Even et al., 2012; Berg et al., 2010). Nature has optimized these microorganisms, enzymes, and metabolic fluxes over billions of years, rendering attempts to improve energy consumption and carbon-fixation efficiency very challenging. Implementing natural or new-to-nature synthetic pathways for C1 assimilation into biotechnologically important microbes such as *E. coli*, which do not naturally grow on C1 compounds, can solve these problems. Assimilation of C1 compounds such as CO_2_, formate, CO, or methanol via their respective assimilation pathways has been receiving increased attention (Bang et al., 2020; Chen et al., 2020; Gassler et al., 2020; Gleizer et al., 2019; Meyer et al., 2018; Schwander et al., 2016). Besides the reductive acetyl-CoA pathway, the synthetic and oxygen-tolerant reductive glycine (rGly) pathway is the most efficient formate assimilation pathway (Bar-Even et al., 2013; Claassens et al., 2022). This pathway was recently successfully engineered in *E. coli* for generation of all biomass from formate and CO_2_ (Figure 1), reaching a doubling time of 9 h and a biomass yield of 2.3 g cell dry weight / mol (CDW/mol) formate (Kim et al., 2020). However, so far bioproduction via synthetic C1-assimilation pathways has not been reported.

**Figure 1.**
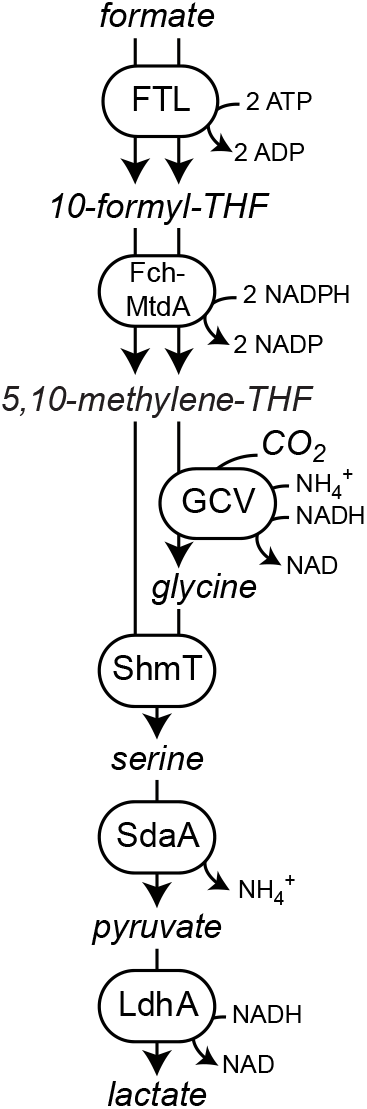
Reductive glycine pathway as operating in the formatotrophic *E. coli* strain. Displayed is formate and CO_2_ conversion to lactate as a final bioproduction product.

In this work, growth performance of the formatotrophic *E. coli* strain was improved using an adaptive laboratory evolution approach on formate under 10% CO_2_ atmosphere. Genome sequencing identified mutations that apparently increased the growth performance on formate, and their effects were verified through reverse engineering. Finally, we demonstrate high biomass production in a fed-batch experiment and engineer lactate production from formate by an optimized *E. coli* strain.

## Materials and methods

### Chemicals and reagents

Primers were synthesized by Integrated DNA Technologies (IDT, Leuven, Belgium). PCR reactions were carried out either using Phusion High-Fidelity DNA Polymerase or Dream Taq (Thermo Fisher Scientific, Dreieich, Germany). Restrictions and ligations were performed using FastDigest enzymes and T4 DNA ligase, respectively, all purchased from Thermo Fisher Scientific. Sodium formate was ordered from Sigma-Aldrich (Steinheim, Germany). ^13^CO_2_ was obtained from Cambridge Isotope Laboratories (Andover, MA, USA).

### Bacterial strains

Wild-type *Escherichia coli* strain MG1655 (F^-^ λ^-^ *ilvG*^-^ *rfb*-50 *rph*-1) was used as the host for all genetic modifications. *E. coli* strains DH5α (F^-^ λ^-^ Φ80*lac*ZΔM15 Δ(*lac*ZYA-*arg*F)U169 *deoR, recA1 endA1, hsdR*17(rK^-^ mK^+^) *phoA supE44 thi-1 gyrA96 relA1*) and ST18 (*pro thi hsdR*^+^ Tp^r^ Sm^r^; chromosome::RP4-2 Tc::Mu-Kan::Tn7λ*pir*Δ*hemA*) were used for cloning and conjugation procedures, respectively. A formatotrophic *E. coli* strain equipped with a reductive glycine pathway (K4e) (Kim et al., 2020) was used as base strain for adaptive evolution and reverse engineering. All strains are listed in Table 1.

**Table 1.**
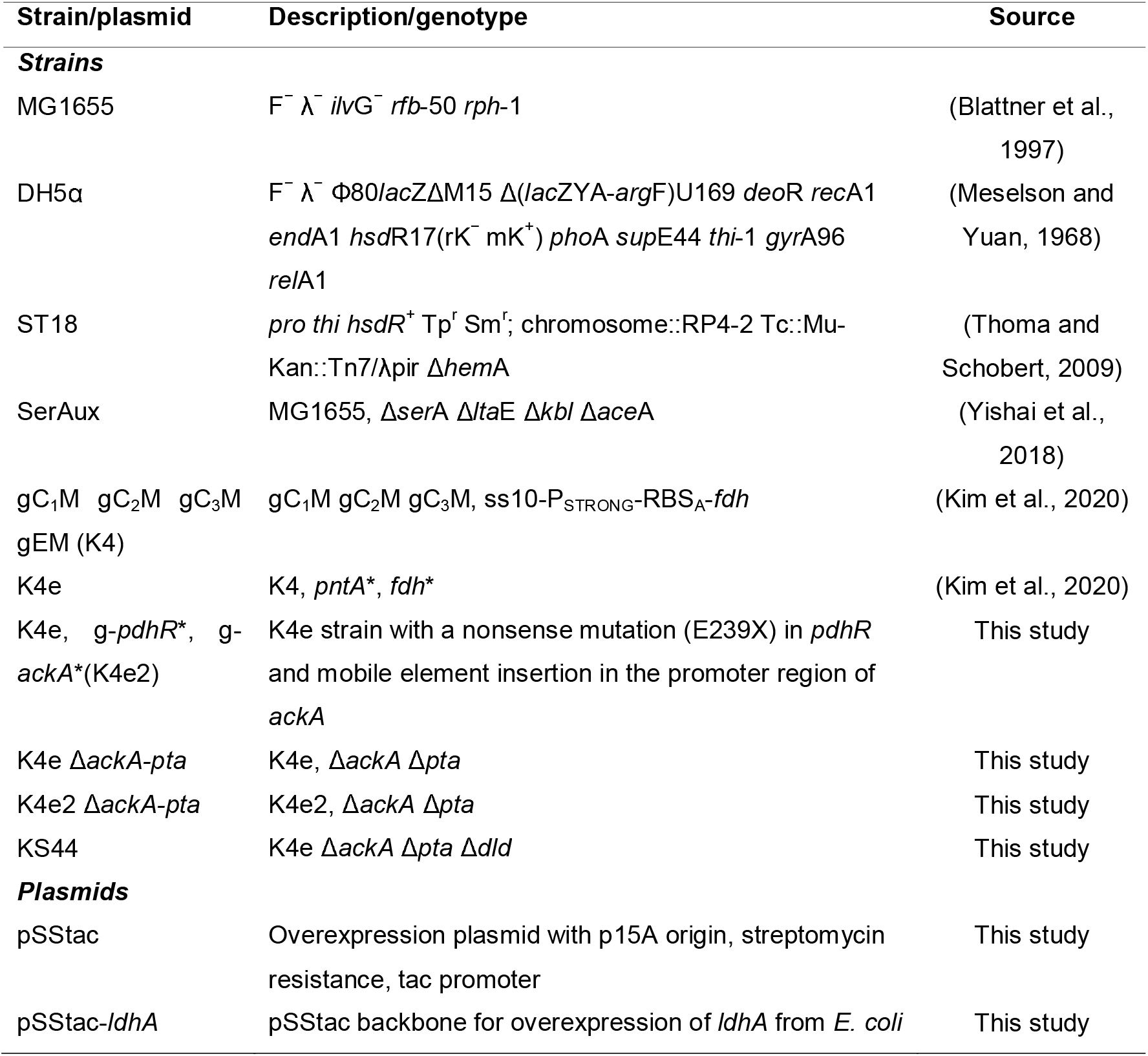
Strains and plasmids used in this study

### Genome engineering

Gene knockouts were introduced in MG1655 by P1 phage transduction (Thomason et al., 2007). Single-gene knockout mutants from the National BioResource Project (NIG, Japan) (Baba et al., 2006) were used as donors of specific mutations. For the recycling of selection markers (as the multiple-gene deletions and integrations required), all antibiotic cassettes integrated into the genome were flanked by FRT (flippase recognition target) sites. Cells were transformed with a flippase recombinase helper plasmid (FLPe, replicating at 30°C; Gene Bridges, Heidelberg, Germany) carrying a gene encoding FLP which recombines at the FRT sites and removes the antibiotic cassette. Elevated temperature (37°C) was subsequently used to cure the cells from the FLPe plasmid.

### Synthetic-operon construction

A gene native to *E. coli*, lactate dehydrogenase (*ldhA*), was prepared via PCR amplification from the *E. coli* MG1655 genome. The PCR product was integrated into a high-copy-number cloning vector pNiv to construct synthetic operons using a method described previously (Zelcbuch et al., 2013). Plasmid-based gene overexpression was achieved by cloning the desired synthetic operon into a pZ vector (15A origin of replication, streptomycin marker) digested with EcoRI and PstI, utilizing T4 DNA ligase. Promoters and ribosome binding sites were used as described previously (Braatsch et al., 2008; Zelcbuch et al., 2013).

### Growth medium and conditions

LB medium (1% NaCl, 0.5% yeast extract, 1% tryptone) was used for strain propagation. Further cultivation was done in M9 minimal media (50 mM Na_2_HPO_4_, 20 mM KH_2_PO_4_, 1 mM NaCl, 20 mM NH_4_Cl, 2 mM MgSO_4_, and 100 μM CaCl_2_) with trace elements (134 μM EDTA, 13 μM FeCl3·6 H_2_O, 6.2 μM ZnCl_2_, 0.76 μM CuCl_2_·2 H_2_O, 0.42 μM CoCl_2_·2 H_2_O, 1.62 μM H_3_BO_3_, 0.081 μM MnCl_2_·4 H_2_O). For the cell-growth test, overnight cultures in LB medium were used to inoculate a pre-culture at an optical density (600 nm, OD_600_) of 0.02 in 4 ml fresh M9 medium containing 10 mM glucose, 1 mM glycine, and 30 mM formate in 10-ml glass test tubes. Cells were then cultivated at 37°C and shaking at 240 rpm. Cell cultures were harvested by centrifugation (18,407 × *g*, 3 min, 4°C), washed twice with fresh M9 medium, and used to inoculate the main culture, conducted aerobically either in a 10-ml glass tube or in Nunc 96-well microplates (Thermo Fisher Scientific) with appropriate carbon sources according to strain and specific experiment. In the microtiter-plate cultivations each well contained 150 μl culture covered with 50 μl mineral oil (Sigma-Aldrich) to avoid evaporation. Growth experiments were conducted (either at 100% air or 90% air / 10% CO_2_) in a BioTek Epoch 2 plate reader (Agilent, Santa Clara, CA, USA) at 37°C. Growth (OD_600_) was measured after a kinetic cycle of 12 shaking steps, which alternated between linear and orbital (1 mm amplitude) and were each 60 s long. OD_600_ values measured in the plate reader were calibrated to represent OD_600_ values in standard cuvettes according to OD_cuvette_ = OD_plate_ / 0.23. Glass-tube cultures were carried out in 4 ml of working volume, at 37°C and shaking at 240 rpm. Volume loss due to evaporation was compensated by adding the appropriate amount of sterile double-distilled water (ddH_2_O) to the culture tubes every two days. All growth experiments were performed in triplicate, and the growth curves shown represent the average.

### Lactate production experiments

Colonies from LB plates for starting liquid cultures in test tubes in LB were incubated at 37°C overnight. All strains grown overnight were washed 3 times with minimal M9 medium. Each strain was then inoculated in test tubes with starting OD_600_ of ~ 0.05. All tubes were incubated in an orbital shaker at 37°C with 10% CO_2_ until the stationary phase in each treatment was reached. Selected cultures were fed with formic acid when each culture reached the stationary phase (OD stopped increasing). Formic acid was added to each tube to increase its concentration by either 30 mM or 60 mM. For each strain, control tubes were left without feeding. Expression of *ldhA* was induced by addition of isopropyl β-d-1-thiogalactopyranoside (IPTG; 1 mM final) together with formic acid. Periodic sampling was performed for measuring the extracellular ions dissolved in the medium by ion chromatography (Dionex ICS 6000 HPAEC, IonPac AS11-HC-4μm Analytical/Capillary Column; Thermo Fisher Scientific). At each sampling point, 200 μl were taken from the cultures and centrifuged at 15,000 rpm for 3 min. The supernatant was then diluted 20 times with ddH_2_O. The diluted sample was centrifuged again at 15,000 rpm for 3 min and transferred to a chromatography vial for the ion chromatography analysis.

### Dry-weight analysis

To determine dry cell weight of *E. coli* grown on formate or methanol, pre-cultures prepared as described above were inoculated to a final OD_600_ of 0.01 into fresh M9 medium containing 90 mM of formate in a 125-ml pyrex Erlenmeyer flask and grown at 37°C with shaking at 240 rpm. Up to 50 ml of cell culture growing in shake flasks were harvested by centrifugation (3,220 × *g*, 20 min). To remove residual medium compounds, cells were washed using three cycles of centrifugation (7,000 × *g*, 5 min) and resuspension in 2 ml ddH_2_O. Cell solutions were transferred to a pre-weighed and pre-dried aluminum dish and dried at 90 °C for 16 h. The weight of the dried cells in the dish was determined and subtracted by the weight of the empty dish. Cell dry weight (CDW) of *E. coli* strains was measured during exponential growth phase (OD_600_ of 0.6–0.8) in the presence of 10% CO_2_ on 90 mM formate.

## Results and discussion

### Adaptive laboratory evolution leads to improved formatotrophic growth characteristics

We previously developed a formatotrophic *E. coli* strain named K4e (Kim et al., 2020). Engineering of this strain was achieved following a modular strategy that included four different modules: (i) a C_1_ module, consisting of formate THF ligase, methenyl-THF cyclohydrolase, and methylene-THF dehydrogenase, all from *Methylobacterium extorquens*, together converting formate into methylene-THF; (ii) a C_2_ module, consisting of the endogenous enzymes of the glycine cleavage system (GCS, GcvT, GcvH, and GcvP), which condenses methylene-THF with CO_2_ and ammonia to give glycine; (iii) a C_3_ module, consisting of serine hydroxymethyltransferase (SHMT) and serine deaminase, together condensing glycine with another methylene-THF to generate serine and finally pyruvate; and (iv) an energy module, which consists of formate dehydrogenase (FDH) from *Pseudomonas sp*. (strain 101), generating reducing power and energy from formate (Kim et al., 2020). After initial growth was observed, the strain’s growth was optimized, reaching performance characteristics of isolated mutants (K4e) of 9 h doubling time and a biomass yield of 2.3 g CDW/mol formate. Subsequent genome sequence analysis and reverse engineering revealed that upregulation mutations in the energy module and the membrane-bound transhydrogenase (a gene product of *pntAB*) supported enhanced growth on formate.

To improve K4e’s growth performance further, we used this strain in an adaptive laboratory evolution (ALE) experiment, selecting for faster growth on formate and CO_2_. The cells were grown in M9 minimal medium containing formate and CO_2_ as the sole carbon sources. We cultivated the K4e strain in test tubes with a formate concentration of 90 mM in a CO_2_ atmosphere set to 10%. Once the turbidity reached an OD_600_ of 1.0, the culture was diluted 1:100 into fresh medium of the same composition to start a new cultivation cycle. While the doubling time gradually decreased over 30 cycles, the final OD_600_ was stagnant for the first 14 cycles (≤ 90 generations). From cycle 15 onwards it appeared that a new mutant became dominant and a stairway-like enhancement in OD_600_ was observed (Figure 2a). To confirm the growth improvement of individuals from the ALE culture, growth of four independent isolates (originating from cycle 26) was analyzed in the plate reader. These independent growth tests conducted with the newly isolated strains, named as ‘K4e2’, confirmed > 40% faster growth of the isolates, reflected by a decrease in doubling time from 9 h to 6.3 h. Strikingly, the isolates also showed a 40% increase in biomass yield, from 2.3 to 3.3 g-CDW/mol formate. Furthermore, increased tolerance toward formate was observed for the newly isolated strain K4e2, which showed accelerated growth as expressed by a reduced doubling time from 8.9 to 6 h and an increase of the final OD_600_ from 0.9 to 1.53 at 150 mM formate (Figure 2c). Moreover, even with 200 mM of formate, the strain grows normally without any significant growth-rate reduction (6.5 h doubling time). Compared to K4e, which showed optimal growth at < 90 mM formate and poor growth at > 150 mM, K4e2 represents a clear improvement. Thus, the isolated strain not only improved biomass productivity, but also in formate tolerance, a feature that is especially beneficial in terms of bioprocess design, since the system can be more robust with strains that exhibit higher formate tolerance.

**Figure 2.**
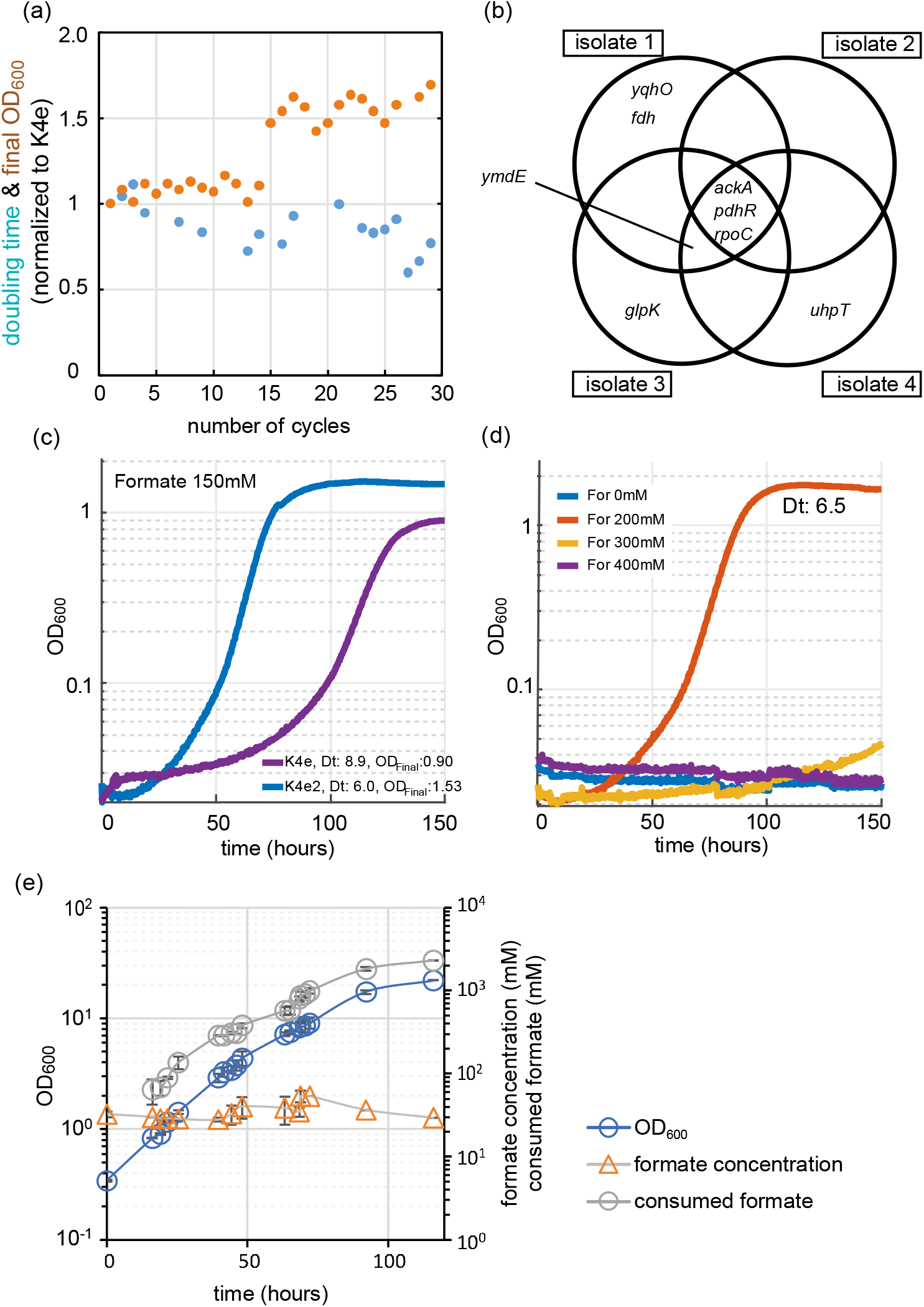
Evolution approaches for enhancing growth on formate. (a) Evolution from K4e to K4e2 via laboratory evolution was conducted in test tubes in M9 minimal medium with 90 mM formate in the presence of 10% CO_2_. Final OD_600_ (orange circle) and doubling times (blue circle) were normalized to K4e. (b) All identified mutated genes obtained after 30 cycles of re-inoculation. Only newly identified mutations are shown compared with parental strain K4e. (c) Growth profile comparison between evolved strains. (d) Formate tolerance test of K4e2. (e) Fed-batch cultivation of strain K4e2 in a bioreactor with pH control. Feeding and pH control were achieved by pumping 10 M of formic acid. List of genes: *pdhR*, pyruvate dehydrogenase complex regulator; *ymdE*, uncharacterized protein; *ackA*, acetate kinase; *yqhO*, biofilm formation related gene; *rpoC*, RNA polymerase subunit beta; *glpK*, glycerol kinase; Experiments (c) and (d) were conducted at 10% CO_2_ in 96-well plates and were performed in triplicates, which displayed identical growth curves (± 5%) and hence were averaged. The corresponding doubling times (Dt) are shown in the figure.

To reveal the genetic changes underlying the growth improvements of the K4e2 isolates, the genomes of all four isolates were sequenced (see supplementary table for all mutations identified). The analysis revealed that all four isolates share three common mutations. The first mutation is a mobile element insertion in the promoter region of *ackA*, which encodes an acetate kinase, responsible for acetate uptake or acetate overflow metabolism in the presence of oxygen (Szenk et al., 2017; Wolfe, 2005). The second mutation is a nonsense mutation (E239X) in *pdhR*, encoding a DNA-binding transcriptional dual regulator. The gene product of *pdhR* represses genes of the pyruvate dehydrogenase complex (PDH) and in the terminal electron transport systems (Ogasawara et al., 2007). As PDH converts pyruvate—a product of formate assimilation via the rGly pathway—to acetyl-CoA, a change in gene expression brought about by a *pdhR* mutation might positively influence growth by decreasing oxidative flux via the TCA cycle. Moreover, the enzyme complexes of PDH and GCS, the key enzyme of the rGly pathway, both contain lipoamide dehydrogenase, the expression of the corresponding gene (*lpd*) is repressed by PdhR (Quail and Guest, 1995). Lastly, a point mutation (A919V) occurs at *rpoC*, which encodes RNA polymerase subunit β’. Here, a direct relevance to carbon and energy metabolism is not obvious (Figure 2b).

### Blocking of acetate overflow metabolism improves formatotrophic growth

Among the mutations found in the K4e2 isolates from the evolution experiment, the mobile element (ME) insertion into the upstream region of *ackA* (acetate kinase) provides important information regarding the formatotrophic growth mode of *E. coli* via the rGly pathway. The ME insertion occurred at the −35 element in the promoter region of *ackA*, which we assume would decrease the level of *ackA* expression. It is not intuitive to consider the occurrence of acetate overflow metabolism in *E. coli* while growing on formate and reaching only very limited final OD_600_ (Basan et al., 2015; Bernal et al., 2016). However, the observed ME insertion suggests that by-product generation might be a limiting factor while growing on formate. Hence, we measured accumulation of metabolites, including acetate, succinate, lactate, and pyruvate during formatotrophic growth. Acetate accumulation was indeed observed from an early growth stage and reached up to 0.2 mM in K4e (Figure 3a). We found that the excreted acetate is re-assimilated as the cell enters the mid-exponential phase. This can be facilitated by either acetate kinase (*ackA*) and phosphotransacetylase (*pta*), consuming one ATP, or acetyl-CoA synthetase (*acs*), which converts acetate to acetyl-CoA while consuming two ATP equivalents (Kumari et al., 1995). In order to prevent K4e from synthesizing acetate, causing loss of carbon and energy, and to mimic the ME insertion found in the K4e2 isolates, both *ackA* and *pta* were deleted, resulting in the K4eΔ*ackA-pta* strain. When K4eΔ*ackA-pta* was cultured using the same conditions, acetate excretion was strongly reduced. To compare the growth performance of K4e and K4eΔ*ackA-pta*, two different formate concentrations were used. With 80 mM formate, both strains show similar growth patterns with almost identical doubling times. However, with 150 mM formate, the K4eΔ*ackA-pta* strain displays not only a reduced doubling time of 7.2 h (as compared to 8.5 h of K4e), but also exhibits a 30% increase in the final OD_600_ (Figure 3b). Indeed, we determined the observed biomass yield of K4eΔ*ackA-pta* to be 3.1 g CDW/mol formate, close to that of K4e2 (3.3 g CDW/mol formate). Moreover, the apparent lag phase of K4e on 150 mM formate was not present in K4eΔ*ackA-pta*, thus the K4eΔ*ackA-pta* strain grew within 80 h to the stationary phase, while the parental strain required more than 140 h to reach the stationary phase. It is unclear how blocking of the acetate overflow metabolism leads to increased formate tolerance. It is conceivable that the complete oxidation of acetyl-CoA via the tricarboxylic cycle provides additional energy to deal with toxicity exerted by formate. Thus, abolishing acetate biosynthesis in K4e is apparently helpful for growth on formate.

**Figure 3.**
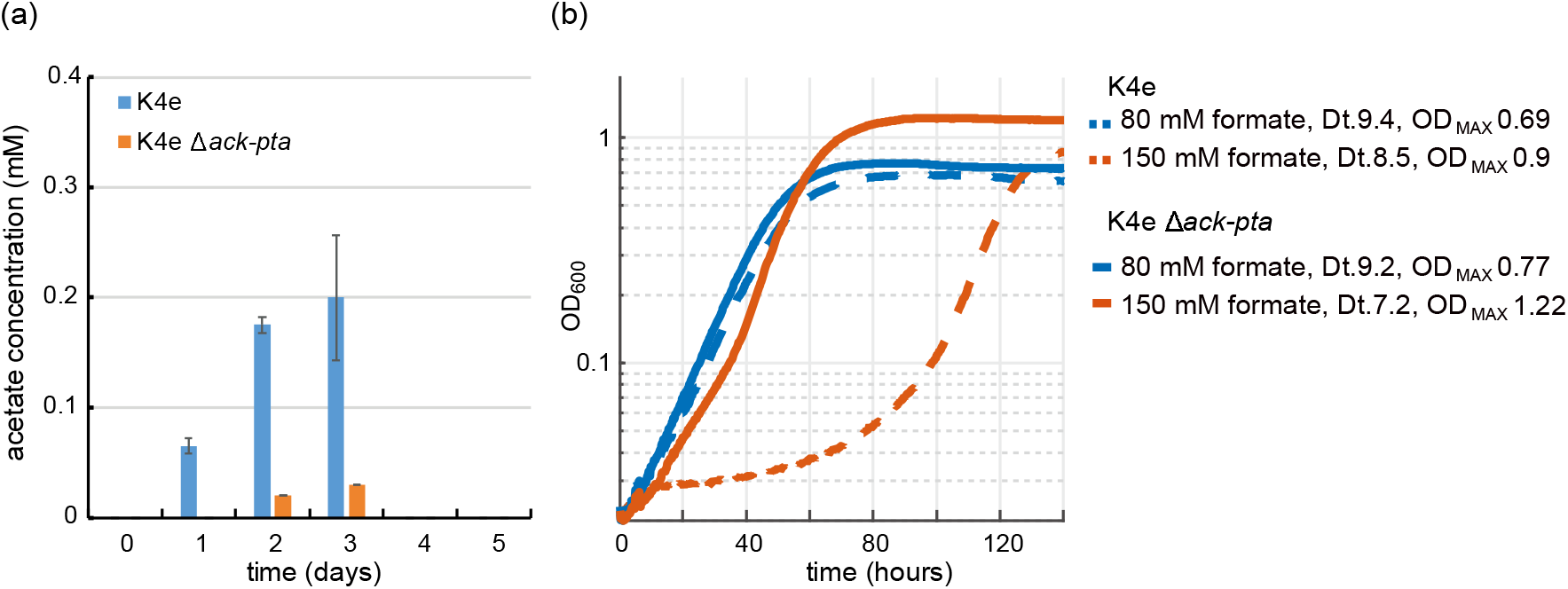
Effect of acetate overflow metabolism on the growth of K4e. (a) Acetate accumulation of K4e in test tubes with 90 mM initial formate. Only acetate was excreted in K4e and re-assimilated as the cell growth enters exponential phase. Strongly reduced acetate production was observed with K4e Δ*ackA-pta*. (b) Growth profile of K4e and K4eΔ*ackA-pta* on 80 and 150 mM formate in 96-well plate experiments performed in triplicates, which displayed identical growth curves (±5%). Deletion of *ack-pta* resulted in increasing final OD_600_ and high formate tolerance. Dt, doubling time.

To further investigate if a complete deletion of *ackA* and *pta* in the evolved K4e2 strain would positively influence the strain’s growth performance, we deleted both genes in the K4e2 strain, yielding K4e2Δ*ackA-pta*. A direct comparison of growth of K4e2 to K4e2Δ*ackA-pta* revealed no difference in terms of doubling times and final OD_600_ when strains grew with 150 or 240 mM formate (Figure 4). However, K4e2Δ*ackA-pta* clearly displays a reduced lag phase before the onset of exponential growth, suggesting that the complete deletions allow the strain to use the formate more efficiently. However, this is not reflected in the observed biomass yield of 3.4 g CDW/mol formate, which is virtually identical to the biomass yield of the parental strain K4e2. However, the biomass yields achieved by K4e2 exceeds the reported average biomass yields of microorganisms naturally growing on formate via the Calvin–Benson–Basham cycle (3.2 g CDW/mol formate) (Claassens et al., 2019).

**Figure 4:**
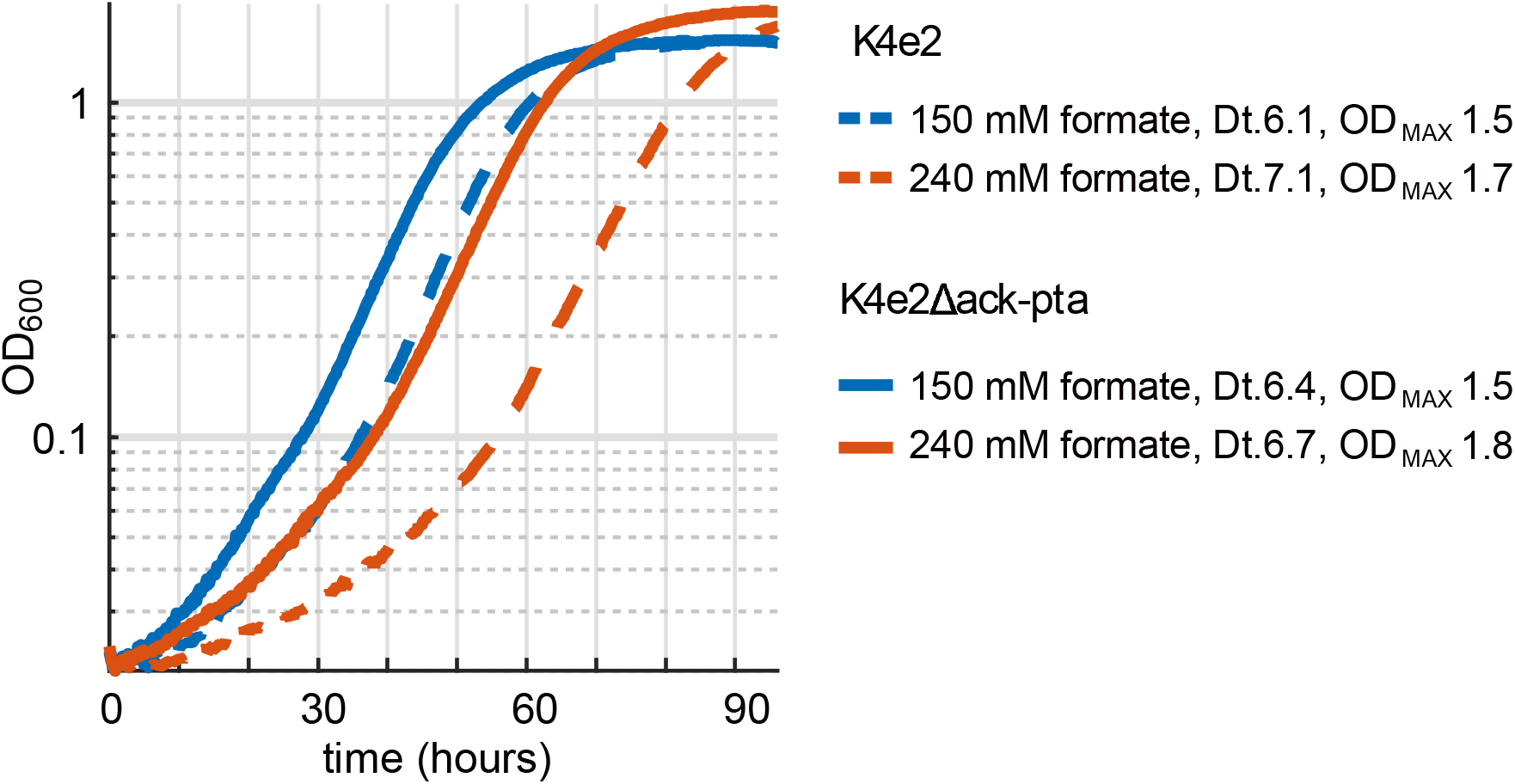
Deletion of *ack-pta* in K4e2 further optimizes formatotrophic growth.

### Fed-batch cultivation for high biomass generation

In order to characterize the strain’s potential for large-scale production we conducted fed-batch cultivation in a 1-l stirred-tank bioreactor, growing strain K4e2 in M9 medium with 30 mM formate at pH 7. Control of pH and feeding of formate was done using 10 M formic acid. Starting from an inoculation OD_600_ of 0.34, the strain reached a final OD_600_ of 22 within 116 h (corresponding to 6 doublings) of incubation with a growth rate of 0.048 h^-1^, corresponding to a doubling time of 14.4 h. The total biomass produced corresponded to 8 g CDW/l (Figure 2e). With a total consumption of 2.289 mol/l formate, the observed biomass yield was determined to be 3.5 g CDW/mol formate, which is consistent with the values derived from batch cultivations. The achieved cell density and growth velocity largely exceeds those previously reported for engineered formatotrophic *E. coli* (Bang et al., 2020). This highlights the potential of the engineered strain for use in industrial bioprocesses, where high cell densities are often required to achieve economic feasibility. However, some improvement with respect to biomass yield and doubling time is still possible, especially when comparing to the reported maximal theoretical biomass yields of ~ 5 g CDW/mol formate (Bar-Even et al., 2013; Cotton et al., 2020). We thus set out to further improve our strains by analyzing and making use of mutations that accumulated during the adaptive evolution.

### Formatotrophic lactate production in *E. coli*

Lactate was selected as a proxy chemical to show the potential of formatotrophic bioproduction. Lactate is an important chemical used in the food and chemical industries as it has a hydroxyl and a carboxyl functional group and can undergo self-esterification to form poly-lactic acid (PLA), a well-known polymer for producing bio-plastic (Maki-Arvela et al., 2014). Lactate can be generated by a reaction catalyzed by lactate dehydrogenase (*ldhA*), which oxidizes NADH using pyruvate as an electron acceptor. In order to prevent *E. coli* from re-assimilating the lactate, quinone dependent D-lactate dehydrogenase (*dld*) was deleted, generating K4eΔ*ackA-pta*Δ*dld*, named KS44. To achieve formatotrophic lactate production, the KS44 strain was transformed with the *ldhA* gene cloned into an IPTG-inducible expression cassette in plasmid pSStac. The final strain with *ldhA* overexpression was named KS46. To test for IPTG-inducible lactate production from formate, we applied a two-phased fed-batch strategy. The growth phase was started in 90 mM formate and continued until the cells entered the stationary phase. The production phase was initiated by adding 1 mM IPTG and 60 mM of formic acid to the culture (Figure 5a). During the growth phase, the pH of the culture increases due to formate uptake into the cell, either via a proton symport mechanism or in the form of free formic acid (Wang et al., 2009; Wei et al., 2011; Wiechert and Beitz, 2017). Assimilation of 90 mM formate increased the culture pH from 6.9 to 7.8 (Figure 5b). Addition of 60 mM formic acid decreased the culture’s pH back from 7.8 to 6.9. Along with the two-phased fed-batch strategy, a normal batch culture was also cultivated for comparison. However, we were able to detect lactate production only in the fed-batch cultivation (1.2 mM; Figure 5c), corresponding to almost 10% of the maximal theoretical yield (Cotton et al., 2020).

**Figure 5.**
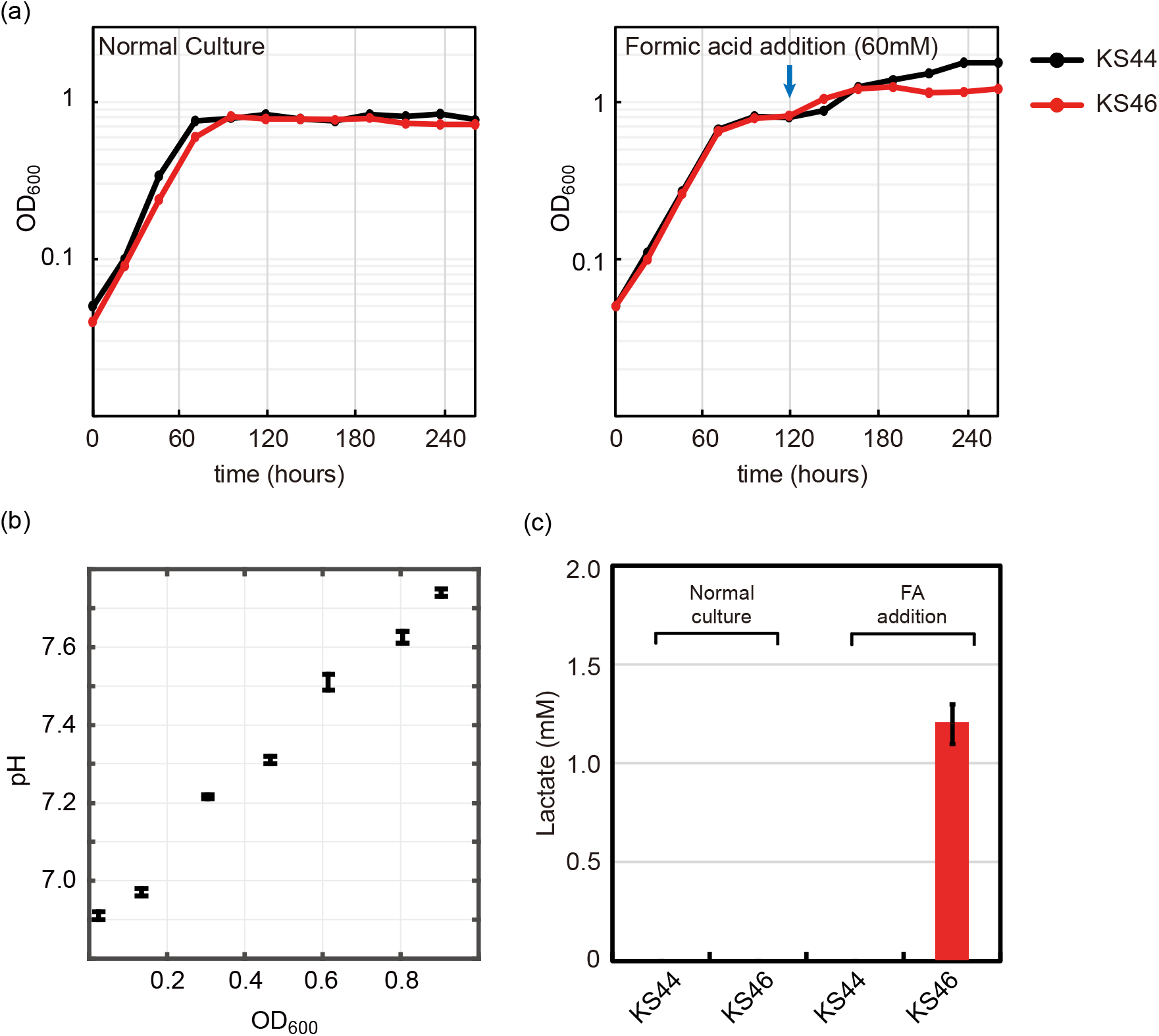
Lactate production from formate with formatotrophic *E. coli*. (a) Engineered *E. coli* strain was cultured in minimal medium using 90 mM formate and 10% CO_2_ as carbon sources. Two different cultivation methods were tested: normal batch mode and fed-batch mode with the addition of 60 mM formic acid (FA) at the indicated time point. (b) pH profile during growth on formate. Cell growth on formate directly correlates with increased medium pH due to the accumulation of OH^-^. (c) Lactate production was observed only with the strain in the fed-batch mode along with *ldhA* overexpression (n = 4, lactate measurement was conducted at the end of cultivation).

## Conclusion

This study demonstrates that a previously engineered formatotrophic *E. coli* strain equipped with the rGly pathway can be optimized for the production of value-added chemicals such as lactate. Adaptive laboratory evolution conducted with CO_2_ and formate as carbon sources yielded a strain with increased biomass yield, shorter doubling time, and the ability to grow to high cellular densities. Interestingly, formate tolerance was increased as well, allowing growth with 200 mM formate, while the parent formatotrophic strain K4e did not grow at such formate concentrations. Subsequent genome sequencing revealed that avoidance of acetate production is one of the key factors for improved growth. By-product analysis of K4e showed that this strain indeed generates acetate during growth on formate and re-assimilates excreted acetate at the late stage of the growth phase. When acetate kinase and phosphate acetyltransferase were deleted from K4e, a similar growth-profile compared to K4e2 was observed, especially with high formate concentration. While the maximal cell density previously reported was 3.5 g CDW/l and the biomass yield 2.5 g CDW/mol formate (Bang et al., 2020), the newly evolved K4e2 strain exceeded those by reaching a cell density of 8 g CDW/l and a biomass yield of 3.4 g CWD/mol formate. This finding not only exemplifies the utility of adaptive laboratory evolution but also constitutes a further step towards the industrial use of the synthetic rGly pathway. This sustainable approach to bacterial biomass production can directly find application in areas like single-cell protein or feed production. However, even more urgent but more challenging is the development of sustainable processes for value-added chemicals to provide alternatives to petroleum-based sources.

In order to achieve biological transformation of formate to lactate, inducible lactate dehydrogenase was implemented and the lactate assimilating quinone-dependent D-lactate dehydrogenase was deleted. When formate/formic-acid fed-batch cultivation was carried out with this strain, production of lactate was observed. We thus showed that our formatotrophic *E. coli* strain, which utilizes formate as energy and carbon source through the rGly pathway, can be further optimized, in this case by prevention of the wasteful acetate formation, and can be applied for the microbial conversion to a chemical of interest. Finally, further strain engineering to increase flux towards lactate and the establishment of an optimized bioprocess will unlock the full potential of the reductive glycine pathway and hence help paving the way towards a C1-bioeconomy.

## Acknowledgements

The authors thank Enrico Orsi and Hezi Tenenboim for critically reading of the manuscript. This work was funded by the Max Planck Society and by the European Union’s Horizon 2020 research and innovation programme under grant agreement No. 763911 (Project eForFuel).

## Conflict of interest

F. K. is cofounder of b.fab, aiming on commercialization of formate-based microbial bioproduction.

