## Supplementary Table for "Optimizing *E. coli* as a formatotrophic platform for bioproduction via the reductive glycine pathway"

Supplementary Table 1. All mutations in K4e2 isolates compare to K4e

| **Mutation** | **Type** | **Annotation** | **Affected enzyme (gene)** |
| --- | --- | --- | --- |
| Point mutation | CDS | E239X (G🡪T) | pyruvate dehydrogenase complex regulator (*pdhR*) |
| Point mutation | CDS | A919V (C🡪T) | RNA polymerase subunit beta (*rpoC*) |
| Mobile element integration | UTR | - | acetate kinase (*ackA*) |
| polymerphism | CDS | - | biofilm formation related gene (*yghO*) |
| Point mutation | UTR | T 🡪 A | *Pseudomonas sp.* formate dehydrogenase (*fdh*) |
| Polymerphism | CDS | - | uncharacterized protein (*ymdE*) |
| Point mutation | CDS | V103I (G🡪A) | glycerol kinase (*glpK*) |
| Point mutation | CDS | Q96X (C🡪T) | hexose-6-phosphate:phosphate antiporter (*uhpT*) |
